# Diverse Microbial Exposure Enhances CD8^+^ T Cell Effector Memory Output and Function

**DOI:** 10.64898/2026.02.02.703037

**Authors:** Claire E. Thefaine, Erin D. Lucas, Katharine E. Block, Mark Pierson, Emma M. Dehm, Matthew A. Huggins, Odhrán W. Casey, David Zemmour, Stephen C. Jameson, Sara E. Hamilton, the immgenT Project

**Affiliations:** Center for Immunology, Department of Laboratory Medicine and Pathology, University of Minnesota, Minneapolis, MN; Department of Immunology, Harvard Medical School, Boston, MA; Department of Pathology, University of Chicago, Chicago, IL

## Abstract

Mice with normalized microbial exposure (NME) harbor an immune system that more accurately reflects that of humans compared to mice maintained as specific pathogen-free (SPF). An explanation for the observed alterations in the composition of the T cell compartment in NME mice has not been reported. We compared the T cell landscape in NME versus SPF mice at baseline and after acute LCMV infection. Using the immgenT dataset, we found no unique T cell populations in NME, but the landscape shifted towards activated T cells with increased propensity for effector functions and improved pathogen clearance. CD8^+^ KLRG1^+^ cells (immgenT CD8_cl12) are significantly expanded in NME mice. Their predominance was a result of both increased formation and the conversion of other memory populations to a KLRG1+ phenotype. Thus, NME mice provide insight into a diverse T cell compartment rich with cells previously found to be limited in SPF mice.

## Main

Normalized microbial exposure (NME) mouse models combine the tractable genetics of common laboratory mouse strains with microbial and pathogen exposure that is more similar to that of adult humans^1-3^. Accordingly, NME mice have been increasingly utilized in immunological and translational research^4-7^. Several different NME models show consistent and durable alterations to the entire immune system, including striking shifts in the T cell compartment^1,3^. In the circulation, CD8^+^ central memory T cells (TCM) (CD44^+^, CD62L^+^, CD127^hi^, KLRG1^-^), CD8^+^ effector memory T cells (TEM) (CD44^+^, CD62L^-^, CD127^hi^, KLRG1^-^), and CD8^+^ long-lived effector T cells (LLEC) or similarly defined terminally-differentiated effector memory (tT_EM_)^8^ (CD44^+^, CD62L^-^, CD127^lo^, KLRG1^+^), form following acute infection and provide protection following reinfection^9^. However, in SPF mice TCM display better homeostatic turnover and persistence compared to TEM and LLEC, leading to their dominance of the memory pool long-term^10^. This contrasts with the CD8^+^ memory T cell compartment of adult humans and NME mice, which are predominantly composed of effector memory subsets^1^. This shift in the CD8^+^ T cell memory compartment may partially explain alterations in responses to vaccines and pathogens in NME mice, including more effective clearance of bacterial infection^1,3^ and reduced responses to vaccination^6,11,12^.

Though several studies have demonstrated considerable alterations to the composition of immune cells in NME mice using flow cytometric based assays, little has been done to examine changes to the transcriptome at the single cell level. Additionally, it is unclear to what extent the surface proteins and transcription factors used to identify T cell populations fully capture cellular heterogeneity in the NME T cell compartment. Characterizing T cells in NME mice is further complicated by the animals’ sustained and systemic pro-inflammatory state, which could promote temporary transcriptional changes in T cells that confound their cellular identity. To address these challenges, we performed single-cell RNA sequencing (scRNA-seq) in collaboration with the immgenT consortium. immgenT is an open-source, community-built reference atlas of mouse T cells that integrates scRNA-seq, CITE-seq, and paired TCR profiling across tissues and immune perturbations to define a flexible, reproducible taxonomy and enable any new dataset to be mapped into a harmonized framework.

To characterize the NME mouse T cell compartment at baseline, we sorted T cells (CD3ε^+^) from the spleens of mice housed in NME conditions for 2 weeks (2wk NME) or 2 months (2m NME), (n=4 each) (Supp. Fig. 1a) (IGT44). scRNA-seq data from these cells was integrated as part of the immgenTv1 reference database and was subsequently projected, along with ImmgenT SPF controls, onto the immgenT integrated framework (Fig. 1a, Supp. Fig. 1b). All T cell populations are represented to some degree in SPF mice that have undergone various single infections or other conditions (Fig. 1b). Of the 113 immgenT clusters, 51 were represented in IGT44 SPF samples, 71 were represented in NME samples, and 50 overlapped between the two. None of the clusters represented in only IGT44 SPF or NME conditions represented more than 2% of a given sample and were primarily effector populations present in 2wk NME samples (Supplementary Table 1). There was a considerable shift in the composition of the splenic T cell compartment in 2wk NME mice, many of which were maintained in 2m NME mice (Fig. 1b). Broadly, we observed an increase in the frequency of CD8αβ^+^ T cells at the expense of conventional CD4^+^ T cells in 2wk NME mice, a trend that was augmented in 2m NME mice (Fig. 1c). To further explore changes to conventional CD4^+^ and CD8αβ^+^ T cells in NME mice, we assessed the representation of immgenT subclusters in our dataset. Within the conventional CD4^+^ T cell population, we saw a notable (∼20%) reduction of CD62L^+^ populations, primarily CD4_cl2 and CD4_cl3, in 2wk and 2m NME samples, suggesting an overall loss of naïve and central memory-like T cells (Fig. 1d). Of the CD62L^-^, CD44+ cells, we observed a durable expansion of Tfh and chronically stimulated cells (CD4_cl9, 13, and 14) in NME samples. At two weeks of NME, there was a slight expansion of Th1-like clusters (CD4_cl21, CD4_cl22) but not Th2-(CD4_cl15, CD4_cl16) or Th17-like clusters (CD4_cl17, CD4_cl18), as expected, given the numerous viral encounters mice experience upon NME housing^1,13^. CD4_cl4, interferon-stimulated gene (ISG)-expressing CD62L^+^ cells, were also transiently expanded only at 2wk NME in accordance with previous reports that interferon signaling peaks within two weeks of NME exposure^14^. Within the CD8αβ^+^T cell compartment, we found an expansion of short-lived effector (SLEC)-like clusters (CD8_cl14,15, and 28) in 2wk NME samples which is largely resolved by two months (Fig. 1e). We saw a dramatic loss in the frequency of resting T cells (CD8_cl1) in 2w and 2m NME samples alongside a considerable expansion of CD8_cl12 (Fig. 1e). CD8_cl12 cells express high levels of *Klrg1* and resemble KLRG1^+^ long-lived effector cells (LLEC), a subset of CD8^+^, CD62L^-^, CD44^+^, CD127^lo^ effector memory T cells that have previously been described by our group^10,15,16^. Additionally, we saw shifts in the composition of both Treg and DN clusters in 2wk NME and 2m NME groups (Supp. Fig. 1d). Taken together, NME doesn’t induce new states of T cells but alters the T cell landscape profoundly compared to SPF mice. NME housing promotes a shift in conventional CD4^+^ and CD8αβ^+^ T cells away from naïve and central memory T cells towards effector (memory) populations, in agreement with previously published flow cytometry data from microbially experienced mice^17^ (Fig. 1f, Supp. Fig. 1e).

**Figure 1:**
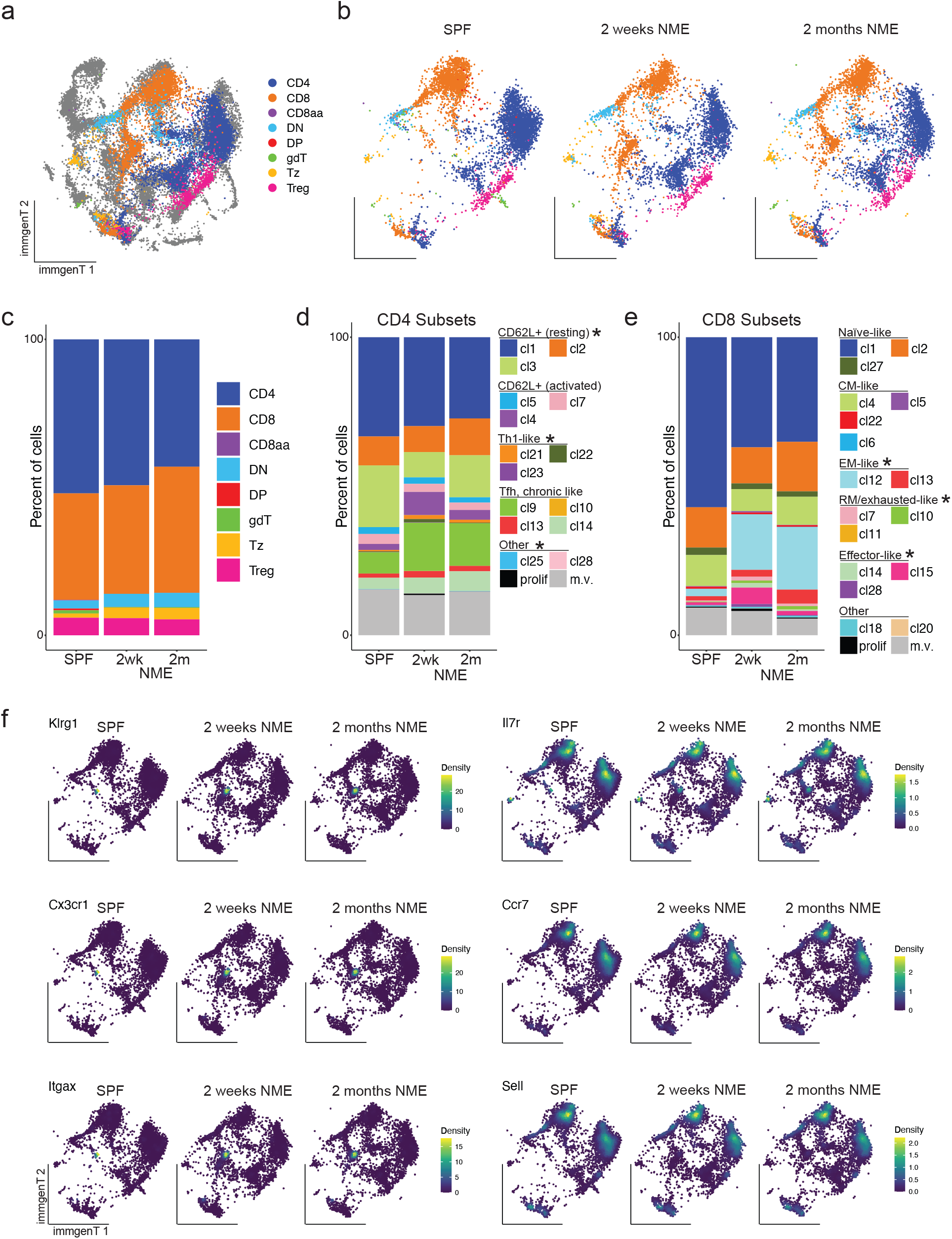
Normalizing microbial exposure leads to maturation of the murine splenic T cell compartment. a) immgenT dimensions of single cell transcriptional data from splenic CD3ϵ^+^ cells from SPF and NME mice after 2 weeks or 2 months of cohousing (n = 4 each), overlaid onto the complete immgenT atlas. b) immgenT dimensions from a split by condition. c) Frequency of major T cell lineages by condition, as annotated by immgenT. d) Frequency of immgenT conventional CD4^+^ T cell subclusters by condition. e) Frequency of immgenT CD8αβ ^+^ T cell subclusters by condition. f) Expression levels of indicated genes in SPF and NME T cells. For d-e, subcluster representation was compared between conditions using an ANOVA. Differentially represented subclusters (FDR < 0.5) are indicated by an asterisk.

In our NME model, SPF laboratory mice are cohoused with mice acquired from local pet stores. As the pathogens harbored by pet store mice differ and transmit to SPF mice at variable levels^18,19^, this model generates animals with unique infection histories, even within a single cage. As a result, the composition of the immune system between NME mice can vary widely. We thus sought to explore compositional and transcriptional changes to blood cells between SPF (n = 1) and NME (n = 4) mice. We selected mice with variable serological profiles (Supp. Fig. 2a) and frequencies of CD8^+^ T cell with an activated phenotype, as determined by CD44 and CD62L expression in the blood. Resulting scRNA-seq data (Fig. 2a) for each sample were integrated to allow for cross-sample comparison (Fig. 2b, Supp. Fig. 2b). Myeloid cells, particularly neutrophils, were observed at higher frequencies in NME mice (Fig. 2c). This increase in circulating neutrophils is closer to the frequencies of neutrophils observed in adult human blood^20^. We observed a slight increase in the expression of ISGs, as well as other activation markers, when comparing neutrophils from NME1 (the least activated sample) and the SPF sample (Supp. Fig. 2c), a trend which was heightened in NME4 (the most activated sample). While CD8^+^ T cells between SPF and NME samples appeared to be more similar than other immune cells, we also observed an increase in ISG expression in NME mouse CD8^+^ T cells, as well as increased expression of *Ccl5* and *Gzma* (Fig. 2d). To further characterize the T cell compartment in NME mice over the spectrum of CD8^+^ T cell activation, we used immgenT Reference-Based Integration (T-RBI). T-RBI enables the integration of T cell query datasets into the immgenT framework: immgenT dimensional reductions and clusters. As expected, 99.9% of cells identified as non-T cells did not map to the immgenT atlas (Fig 2e).

**Figure 2:**
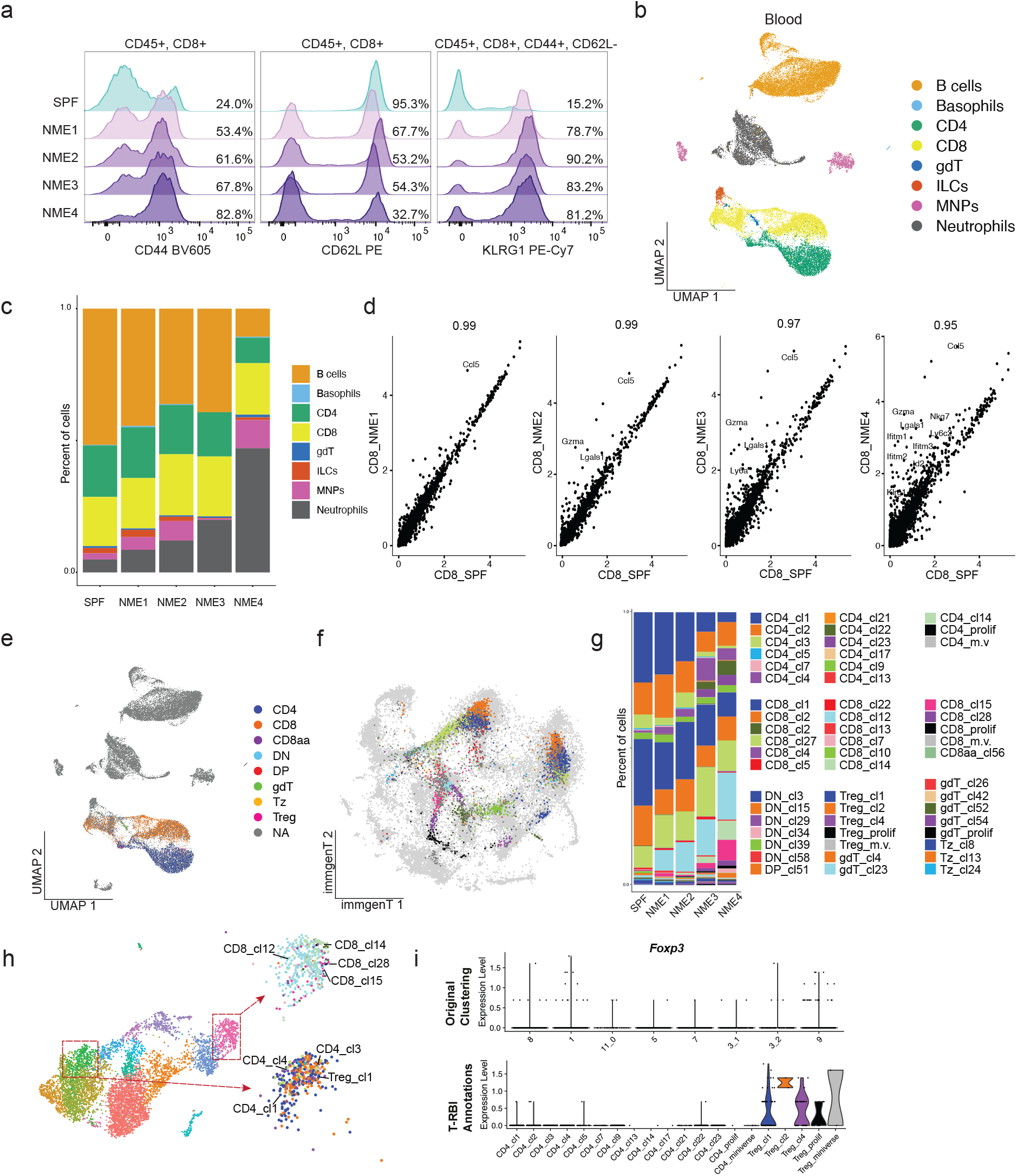
The composition of WBCs vary between NME mice while retaining consistent trends. a) Histograms and quantification of the frequency of CD44, CD62L and KLRG1 expression on indicated cells for each mouse (SPF n = 1, NME n = 4) as assessed by flow cytometry of peripheral blood. b) Integrated UMAP of single cell transcriptional data from blood cells from mice in a. c) Frequency of cells identified in b in each sample. d) Comparison of aggregated CD8 T cell gene expression between SPF and NME samples. e) Integrated UMAP as in b overlaid with immgenT Reference-Based Integration (T-RBI) annotations. f) T-RBI of data in e, in immgenT dimensions and cell annotation. g) Frequency of immgenT subclusters in each sample. h) UMAP of CD3ϵ^+^ blood cells colored by Seurat-generated clusters (left), compared to T-RBI annotations for 2 example clusters (right). i) *Foxp3* expression in CD4^+^ clusters by Seurat-generated clusters (top) and of T-RBI annotations (bottom).

We were interested in comparing our in-house annotations to immgenT subclusters. To this end, we subsetted the data on T cell clusters, reclustered, and reannotated the resulting populations. While the *Seurat* clustering performed under our own analysis was able to distinguish between CD4^+^, CD8^+^, and γδT cells, and the level of differentiation of these cells to some degree, it proved difficult to confidently identify subsets within these lineages, a described limitation of utilizing transcriptional data to separate T cells^21^. Leveraging the immgenT cluster identities determined after T-RBI, we were able to validate our own interpretations of the data and observe more subtle compositional changes in T cell subsets (Fig. 2f,g). The majority (93% of CD4s and 96% of CD8s, Supplementary Table 2) of cells from these clusters aligned with the lineage-level immgenT annotation (e.g. CD8, CD4, Treg). But Seurat clusters were coarse and represented by a diverse range of immgenT subclusters (Fig. 2h). Within the CD8^+^ lineage, the SLEC-(CD8_cl14,15,28) and LLEC-like (CD8_cl12) cells were annotated and separated in the immgenT dimensions, and similar to what we observed in Figure 1, we saw a consistent expansion of CD8_cl12 (LLEC) in each NME mouse (Fig. 2g,h). Interestingly, Tregs, which were not identified in our original analysis, were annotated by T-RBI. In fact, these Tregs were mixed with conventional CD4^+^ T cells in the original UMAP, and a few *Foxp3*^+^ cells appeared in Seurat clusters 8, 1, 3_2, and 9. However, they were clearly separated in the immgenT dimensions, and Treg_cl1, 2, and 4 were distinguished (Fig. 2f,I, Supp. Fig. 2d). Thus, T-RBI allowed us to annotate rare cells that do not form separate clusters within our dataset and identify the diverse T cell responses more accurately.

In summary, while the differentiation of the CD4^+^ and CD8^+^ compartments in these samples appears to increase with the frequency of activated CD8^+^ T cells, the expansion of CD8_cl12/LLEC is apparent even in our least activated sample, indicating that these changes in T cell identity cannot be attributed to a specific pathogen. Furthermore, we did not observe unique-to-NME immgenT clusters in any sample.

As both the host environment and the particular microbial insult influence the differentiation of T cells during primary infection, we sought to understand how the altered, more inflammatory NME environment^3^ affects the differentiation of antigen-specific CD8^+^ T cells during a novel LCMV infection. Thus, we infected age-matched SPF and NME mice with LCMV-Armstrong. Thirty days post-infection (d.p.i.), we sorted CD8^+^, LCMV tetramer^+^ (gp33/H2-D^b+^, np396/H2-D^b+^, or gp276/H2-D^b+^) splenocytes for scRNA-seq as part of the immgenT reference dataset (Fig. 3a) and highlighted our data onto the immgenT CD8αβ atlas (Fig. 3b). Similar to the blood analysis, CD8_cl12 was particularly expanded among LCMV-specific cells in NME mice (Fig. 3c) again at the expense of CD62L^+^ clusters, primarily CD62L^+^ CD44^-^ CD8_cl4 (Fig 3b-d). We applied module scores based on previously described transcriptional signatures of TCM and LLEC^8,10^ to the combined data and found a high degree of overlap between LLEC signature-enriched clusters and NME cells, and conversely, TCM signature-enriched clusters and SPF cells (Fig. 3e, Supp. Fig. 3b).

**Figure 3:**
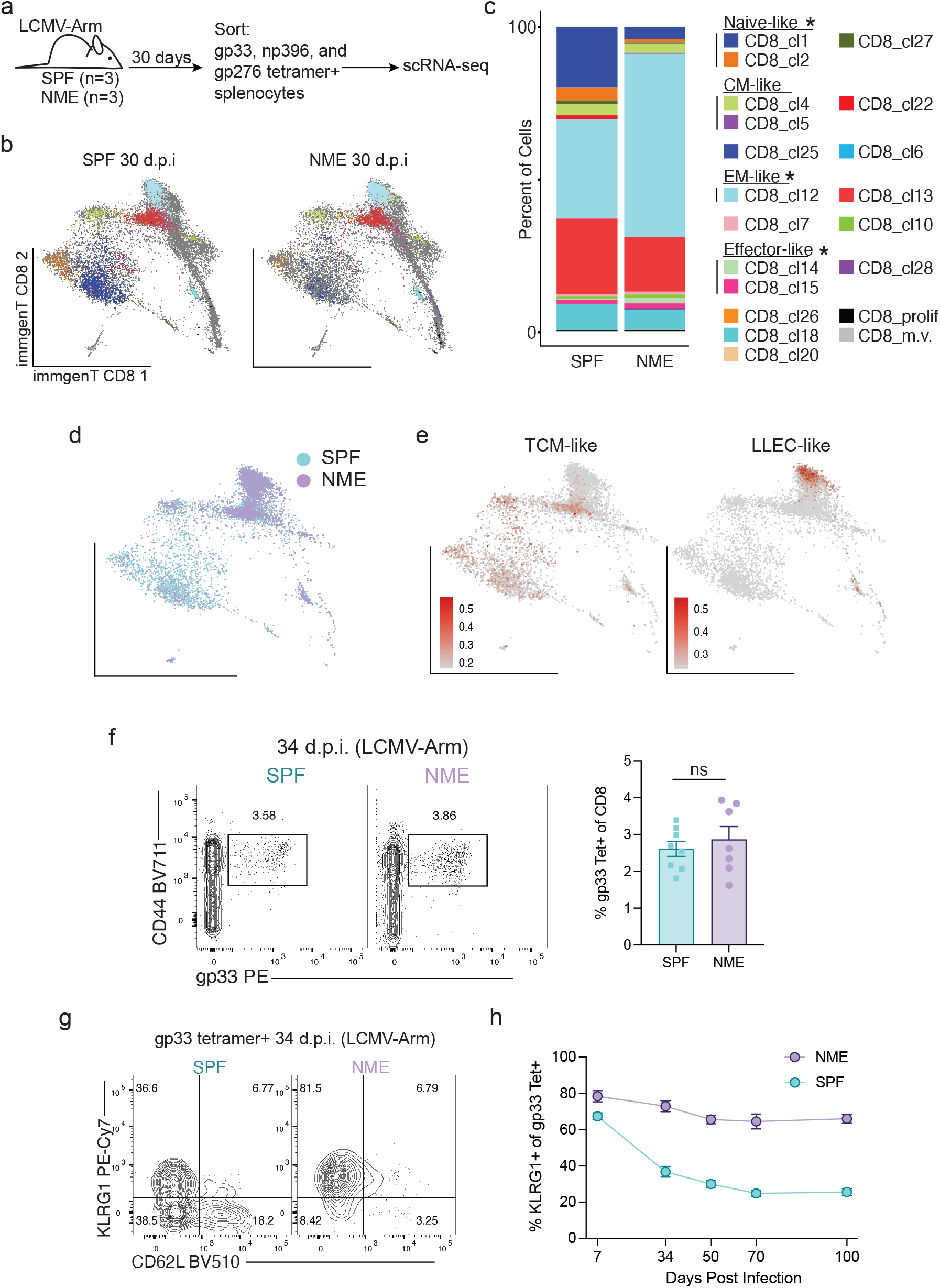
LCMV antigen-specific CD8 T cells in NME mice are more differentiated following a novel infection. a) Experimental outline for b-e. b) immgenT CD8αβ dimensions of single cell transcriptional data from sorted gp33 H2-Db^+^, np396 H2-Db^+^ or gp276 H2-Db^+^ CD8^+^ splenocytes from SPF and NME samples (n = 3 each) 30 d.p.i with LCMV-Arm, overlaid onto the complete immgenT CD8αβ atlas. c) Frequency of immgenT CD8αβ ^+^ T cell subclusters by condition. d) immgenT CD8αβ dimensions as in a, colored by cells from SPF (blue) and NME (purple) samples. e) Module scores of TCM-like or LLEC-like transcriptional signatures. f) Representative flow cytometry plots and quantified frequency of gp33 tetramer^+^, CD8^+^ cells in blood of SPF and NME mice 34 d.p.i with LCMV-Arm. g) Representative flow cytometry plots of KLRG1 and CD62L expression on gp33 tetramer^+^ CD8^+^ T cells from g. h) Frequency of gp33 tetramer^+^ CD8^+^ T cells in the blood of mice from g over time. Data in f-h is combined from two independent experiments (n = 3-4 mice per group in each experiment). For c, subcluster representation was compared between conditions using a moderated t-test. Differentially represented subclusters (FDR < 0.5) are indicated by an asterisk. For f an unpaired, two-tailed Student’s t tests was used yielding a P-value of 0.5167.

To further explore these findings and determine the durability of LLEC in NME mice, we infected SPF and NME mice with LCMV-Armstrong and tracked circulating gp33 tetramer^+^ cells over time. Though we found similar frequencies of gp33 tetramer^+^ cells in the blood of SPF and NME mice (Fig. 3f), there was a striking increase in the proportion of LLEC in NME mice, with around 80% of the tetramer^+^ cells expressing KLRG1 at 34 d.p.i. (Fig 3g), attributable to a loss in the frequency of CD62L^+^ TCM (Supp. Fig. 4a). Intriguingly, NME mice maintain a high frequency of LLEC at 100 d.p.i., whereas LLEC in SPF mice decline over time (Fig. 3h). This trend is not specific to LCMV, as it was replicated in OVA-specific endogenous T cells primed in mice infected with vesicular stomatitis virus-expressing OVA (VSV-OVA) (Supp. Fig. 4b). As the NME method introduces mice to microbial exposures primarily during the early days of cohousing, we were curious to see if mice in prolonged NME conditions still exhibited an increase in LLEC formation and persistence. Even after 150-200 days of NME housing, mice subsequently infected with LCMV-Armstrong still exhibited a higher proportion of LLEC than their age-matched SPF counterparts (Supp. Fig. 4c).

Though the composition of the CD8^+^ memory pool is significantly more differentiated in NME mice, all clusters present in NME mice were also present in SPF mice after accounting for stochasticity (Supplementary Table 3). If the immgenT clusters are truly identifying T cell subsets and not states, then within-cluster variation between conditions should be minimal. Indeed, when we queried the top five represented immgenT defined clusters in our samples (CD8_cl1,4,12,13, and 18), the only differentially expressed genes (DEGs) (expressed in > 25% of either SPF or NME cells, > 1 avg log2FC) were *Junb* (CD8_cl1,4,12, and 13) and *Itga4* (CD8_cl1) (Supplementary Table 4). We were particularly interested in CD8_cl12 given their overwhelming expansion in NME mice. After lowering the log2FC threshold to 0.5, DEG analysis revealed a slight upregulation of *Gzma* in NME CD8_cl12 cells (Fig. 4a). We also found that following *ex vivo* stimulation with gp33 peptide (Fig. 4b,c) a greater frequency of LLEC from NME mice produced IFNγ, which aligns with the increased presence of this subset in NME mice. Intriguingly, when we examined the ability of IFNγ^+^ LLEC to also produce TNFα we found an increase in the frequency of double positive cells (Fig. 4c), suggesting the LLEC responding to peptide are more polyfunctional in NME mice. We observed a similar increase in polyfunctionality of LLEC following PMA/Ionomycin stim (Fig. 4d,e). We next asked if LLEC from NME mice exhibit intrinsic increases in function. We transferred P14 cells into SPF or NME mice and infected them with LCMV-Armstrong. At 30 d.p.i we sorted LLEC P14 T cells generated in SPF or NME mice and transferred purified cells into new, naïve SPF hosts. The next day, we challenged recipient mice with *Listeria monocytogenes*-expressing gp33 (*Lm*-gp33) and quantified colony forming units (CFU) three days later (Fig. 4f). At this early timepoint, we saw a significant reduction in CFU in the spleens of mice that received NME LLECs compared to mice receiving SPF LLECs (Fig. 4g). Altogether, these results suggest that LLEC formed in NME mice have increased effector functionality that can lead to more rapid control of pathogen infection.

**Figure 4:**
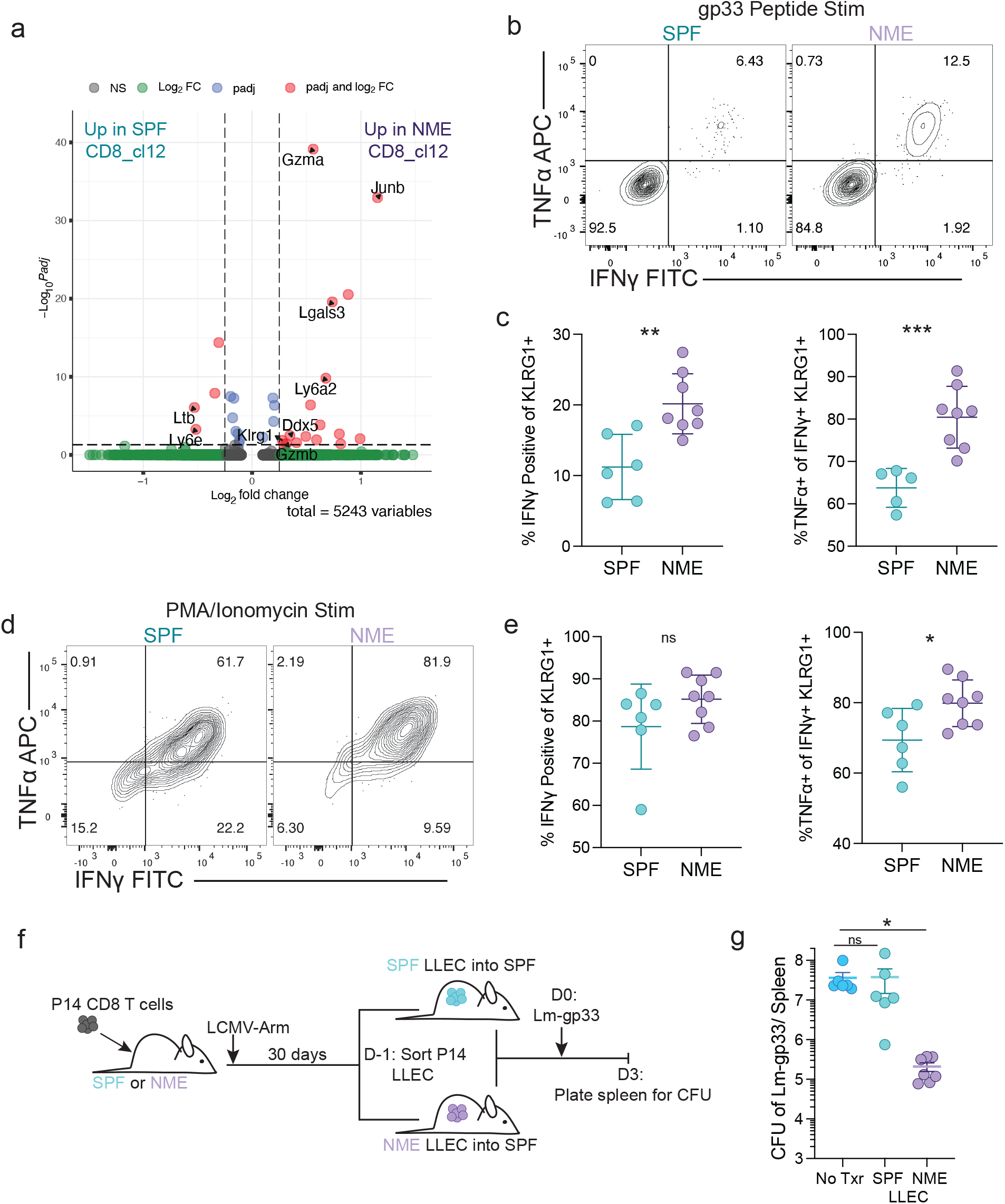
LLEC from NME mice exhibit increased effector functionality. A) Volcano plot of differentially expressed genes between SPF and NME mice within CD8_cl12 (LLEC) from LCMV specific T cells at 30 d.p.i.; data from Fig. 3. b) Representative flow cytometry plots of IFNγ and TNFα expression on KLRG1^+^, CD44^+^, CD8^+^ T cells in SPF and NME mice following stimulation with gp33 peptide. c) Frequency of IFNγ^+^ and TNFα^+^ cells from B. d) Representative flow cytometry plots of IFNγ and TNFα expression on KLRG1^+^, CD44^+^, CD8^+^ T cells in SPF and NME mice following stimulation with PMA and ionomycin. e) Frequency of IFNγ^+^ and TNFα^+^ cells from e. f) Experimental outline for g. g) Quantification of CFU/spleen in SPF and NME mice 3 d.p.i. with *Lm*-gp33. For b-g, data shown is combined from two independent experiments, at 30 d.p.i. (n = 3-4 mice per group). For c and e unpaired, two-tailed Student’s t tests were used, and for g a 1-way Anova followed by Tukey’s multiple comparisons test was used.

We next explored the factors that support the long-term maintenance of KLRG1^+^ T cells in NME mice. In SPF mice, we have previously shown that LLECs exhibit decreased homeostatic proliferation compared to TCM and TEM, contributing to their attrition over time^10^. Given the durable increase in serum cytokines we observe in NME mice^3^ (Supp. Fig. 5a), we wondered if LLEC division is increased in NME mice. Surprisingly, BrdU incorporation was equivalent for gp33 tetramer^+^ LLEC in SPF and NME mice 30 d.p.i. with LCMV-Armstrong, though substantially lower than BrdU labeling in TCM in either group (Supp. Fig. 5b,c). Other groups have shown that some responding CD8^+^, KLRG1^+^ effector T cells downregulate KLRG1 and differentiate into KLRG1^-^ memory subsets in SPF mice^22^. As SLECs (short-lived effector cells, KLRG1^+^, CD127^-^) are induced in part by signals from pro-inflammatory cytokines^23^, we wondered if SLEC formation and maintained expression of KLRG1 would be increased in NME mice. To test this, we sorted gp33 tetramer^+^ SLECs and MPECs (memory-precursor effector cells, KLRG1^-^, CD127^+^) from SPF mice 7 d.p.i. with LCMV-Armstrong and transferred the cells into either SPF or NME hosts (Fig. 5a, Supp. Fig. 5d). Indeed, the transfer of SLECs into NME mice led to stronger formation of LLECs, with ∼80% of transferred cells still present at day 26 expressing KLRG1 in NME mice compared to ∼40% in SPF mice (Fig. 5b,c). Furthermore, ∼40% of MPECs transferred into NME hosts still present at day 26 formed KLRG1^+^ LLECs, as compared to only ∼7% of those transferred into SPF hosts, indicating that the NME environment promotes the conversion of MPECs into KLRG1^+^ SLEC/LLECs (Fig. 5b,c). After establishing the increased formation of LLECs during the effector stage of an immune response in NME mice, we further assessed the influence of NME in the adoption of the LLEC-phenotype by established TCM and TEM cells. We thus immunized SPF mice with gp33-TriVax (peptide, poly(I:C), and agonist anti-CD40 antibody), which induces an overwhelmingly CD8^+^, KLRG1^-^ T cell memory population^24^, and moved these mice to NME housing after memory was established (Fig. 5d). We tracked the gp33 tetramer+ cells in the SPF and NME mice over time, and found that following transfer to NME, the frequency of CD62L^-^, KLRG1^+^ cells increased drastically (Fig. 5e). Together, these data support our hypothesis that the NME host environment promotes the formation of SLECs during the effector stage and also encourages LLEC from other memory populations.

**Figure 5:**
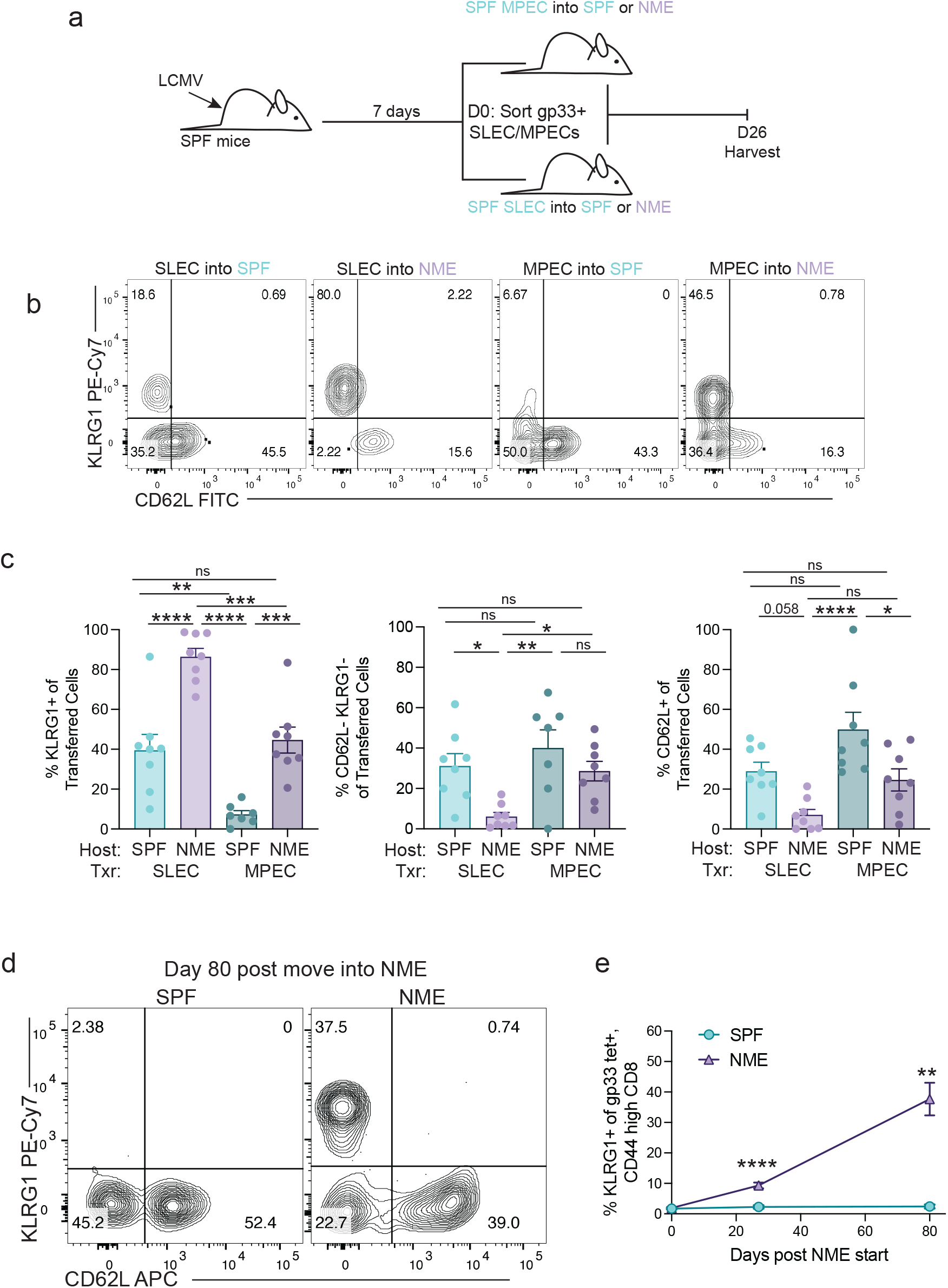
The maintenance of LLECs in NME mice is supported by conversion from other CD8^+^ memory T cell subsets. a) Experimental outline for b and c. b) Representative flow cytometry plots of KLRG1 and CD62L expression on indicated populations. c) Frequency of indicated populations 26 days after adoptive transfer. d) Representative flow cytometry plots of KLRG1 and CD62L expression gp33 tetramer^+^, CD8^+^ T cells in the blood of mice 80 days after NME housing. gp33^+^ memory cells (KLRG1^-^) were induced with gp33-TriVax in SPF mice prior to NME housing. e) Frequency of KLRG1^+^, gp33^+^, CD8^+^T cells in SPF and NME mice over time following NME initiation. Data shown is combined from two independent experiments with 4-5 mice per group. For c a 1-way Anova followed by Tukey’s multiple comparisons test was used, and for e unpaired, two-tailed Student’s t tests were used for each timepoint.

As NME mouse models are a relatively recent development, their purported relevance to the human immune system has been met with healthy skepticism. In our model of NME, adult laboratory mice are rapidly exposed to the micro- and pathobiomes of pet store mice, in contrast to the more gradual microbial exposure experienced by humans, leading to concerns that T cells in these mice may not be physiologically relevant. Conversely, there has been concern that T cells from SPF mice may lack translatability due to the host’s lack of microbial exposure. In this study, we showed that all immgenT reference clusters from NME mice are represented in SPF mice, albeit at different frequencies. In fact, the T cells present in NME mice were also present in numerous experimental contexts included in the immgenT database (see immgenT website https://rstats.immgen.org/immgenT_interactive_maps/CD8.html) with very few transcriptional differences within clusters of the same identity between NME and all other immgenT reference data. Thus, the lack of NME-specific T cells further supports the use of SPF and NME mice to study T cell responses. Given that adult humans have much larger effector memory populations (TEM, TEMRA) than SPF mice^1^, the NME model may more faithfully represent the relative composition of subsets in the human CD8^+^ T cell memory pool.

In line with previous studies, our transcriptional data shows a consistent and durable maturation and expansion of memory cells in the T cell compartment in NME mice. Here, we showed that CD8^+^, KLRG1^+^ LLECs (CD8_cl12) are represented at high frequencies long-term in NME mice. We experimentally determined that KLRG1^+^ SLECs (CD8_cl14) form at higher frequencies in NME hosts and that the NME environment supports the continued expression of KLRG1 on effector cells during the contraction phase, leading to improved LLEC formation. Additionally, we observed KLRG1 expression on formerly KLRG1^-^ memory CD8^+^ T cells after introducing NME, indicating that antigen-independent mechanisms can adjust memory T cell differentiation. Due to the heightened levels of many cytokines and chemokines in the serum of NME mice^25^ (Supp. Fig. 5a), we posit that the inflammatory environment of NME mice promotes the “conversion” of TCM and TEM to LLEC. This data further supports the growing evidence that the T cell compartment is flexible, with established memory cells able to take on different phenotypes in response to environmental triggers^16,26,27^.

By leveraging the immgenT atlas we characterized T cells from the spleen and blood of NME mice, providing insight into the compositional differences between SPF and NME mouse T cell compartments. We further explored the functional consequences of the NME-induced T cell compartment, finding improved LLEC function and pathogen clearance by NME-derived cells. Whether this reflects the modest transcriptional differences between NME and SPF LLEC or distinctions in their epigenetic profiles must await further investigation. Finally, we show that the numeric shifts in T cell populations in NME are likely to be promoted by inflammatory cues rather than only antigenic stimulation. Altogether, this work demonstrates the power of the immgenT atlas as a tool to explore T cell phenotype and function with increased depth and nuance.

## Methods

### Mice

C57BL/6 mice were purchased from the Jackson Laboratory or National Cancer Institute via Charles River. LCMV-DbGP33-specific TCR transgenic P14 mice were fully back-crossed to B6 mice with the introduction of congenic marker CD45.1 for identification. All SPF animals were maintained under specific pathogen–free conditions at the University of Minnesota and all experimental procedures were approved by the Institutional Animal Care and Use Committee. Due to the need to cohouse adult mice in NME, female mice were used in all experiments.

### Pet store mouse cohousing

Pet store mice were purchased from Twin Cities area pet shops and cohoused with SPF mice as previously described^13^. One pet store mouse was cohoused within 1 week of purchase with four 8-week-old laboratory mice in a standard mouse cage. One pet store mouse was introduced per cage and remained in the cage for the duration of the experiment, except for NME4 (Fig. 2) which was cohoused with two pet store mice. Mice were cohoused for two weeks or two months prior to sorting for sequencing. For experiments examining LCMV responses in NME and SPF mice, NME mice were cohoused for at least two months prior to LCMV infection. Serology testing against common mouse pathogens was conducted by Charles River Research Animal Diagnostic Services.

### Viral and Bacterial Infections

Mice were infected with 2e5 PFU of LCMV Armstrong intraperitoneally (i.p.) to generate endogenous LCMV responsive T cells or P14 memory T cells. Mice were infected with 1e6 PFU of VSV-OVA intravenously (i.v.) to generate endogenous ovalbumin specific T cells. *Listeria monocytogenes* expressing gp33 peptide (provided by Hao Shen, University of Pennsylvania) was grown to log phase in tryptic soy broth containing streptomycin (Sigma). Mice were infected with 1e5 CFU as measured by OD600 and confirmed via plating. Bacterial load in the spleen was measured at 3 d.p.i. via homogenization in 0.1% IGEPAL (Sigma) and plating on TSB agar plates.

### Trivax Immunization

Mice were injected i.v. with synthetic gp33 peptide (100ug/mouse, H2N-KAVYNFATM-OH, Biosynth), polyI:C (50ug/mouse, Invitrogen) and agonistic anti-CD40 mAb (100ug/mouse, Bio X Cell).

### Adoptive Transfers

P14 CD8^+^ T cells were magnetically isolated from the spleen of congenic donors. Spleens were passed through a 70µm cell strainer (Corning) and then CD8^+^ T cells were isolated using the MojoSort Mouse CD8^+^ T cell Isolation Kit (BioLegend). Approximately 5e4 purified naïve P14 T cells were transferred i.v. into recipient mice. For transfer of effector cell subsets, spleens were homogenized and stained with fluorescent antibodies. Endogenous gp33 tetramer+ (provided by the NIH tetramer core) SLEC (CD44^hi^, KLRG1^+^, CD127^lo^) and MPEC (CD44^hi^, KLRG1^-^, CD127^hi^) cells were then sorted on a FACSAria II cell sorter (Supp. Fig. 5d). After sorting, purified SLEC (2-4e6 cells/mouse) or MPEC cells (5-7e5 cells/mouse) were transferred into naïve mice.

### BrdU Treatment

For BrdU treatment, mice were injected i.p. with BrdU (2mg/mouse, Invitrogen) for two days, and then given BrdU (0.8mg/mL) in drinking water with 2% sucrose for an additional 12 days. The water was protected from light and changed daily. Following treatment, spleens were harvested and then processed for flow cytometry.

### Cytokine Stimulation

For stimulation with peptide or PMA/ionomycin, approximately 1e6 splenocytes were plated into a 96 well plate. Cells were then resuspended with gp33 peptide (2mg/mL, Biosynth) or PMA (20ng/mL, Sigma) and ionomycin (1μg/mL, Sigma) in media with 1X brefeldin A (Tonbo). Cells were stimulated for 5 hours at 37°C and then stained for flow cytometry.

### Flow Cytometry

Peripheral blood was collected in 10uL of heparin and then red blood cells were lysed with ACK lysis buffer. Spleens were collected and homogenized into a single cell suspension. Cells were then stained for surface antibodies (CD8 BUV395 or APC; CD44 BUV496, BV711 or FITC; CD127 BUV737 or PE; CD45.1 BV421; CX3CR1 BV786; CD62L FITC, PerCP-Cy5.5 or BV510; CD45.2 PerCP-Cy5.5; and KLRG1 PE-Cy7) for 30 min at 4°C. Cells were also stained with Ghost Dye™ Red 780 (Tonbo) to differentiate live and dead cells. We used flow cytometry to define T cell subsets as follows: TCM were defined as CD8^+^, CD44^+^, CD62L^+^, CD127^hi^, KLRG1^-^; TEM were defined as CD8^+^, CD44^+^, CD62L^-^, CD127^hi^, KLRG1^-^; and LLEC were defined as CD8^+^, CD44^+^, CD62L^-^, CD127^lo^, KLRG1^+^. For intracellular cytokine staining, following surface stains, cells were washed and then fixed and permeabilized using the BD Cytofix/Cytoperm Fixation and Perm Solution Kit. Cells were then stained with intracellular antibodies (IFNγ FITC, TNFα APC) overnight 4°C. For BrdU staining, after surface stain, cells were fixed and permed with the Tonbo Transcription Factor Staining Kit and then digested with DNase 1 at 300ug/mL for 55min at 37°C. Following DNA digest, cells were stained with anti-BrdU (FITC) overnight at 4°C.

### Measurement of Circulating Cytokines

Serum cytokines were measured using a custom panel (IL-1β, IL-2, IL-12, IL-4, IL-18, IFN-γ, IL-6, IL-10 and TNF-α) on an xMAP® INTELLIFLEX using software Belysa 1.0.19 via the UMN Cytokine Reference Laboratory.

### Quantification And Statistical Analysis of Flow Cytometry and Cytokine Data

Software Graph-Pad Prism 9.0 was used for statistical analysis. To calculate statistical significance, unpaired Student’s t tests, paired Student’s t tests and 1-way Anova followed by Tukey’s multiple comparisons test for three or more groups were used. For each experiment, there was no significant variation within groups and the variation between groups was similar. All error bars displayed show the mean and standard error of the mean (SEM). Please see figure legends for details of statistics for individual plots. P values were determined with α = 0.05. *P < 0.05, **P < 0.01, ***P < 0.005, and ****P < 0.001.

### Single-cell RNA and Totalseq C-Sequencing - Sample Preparation (immgenT) Flow cytometry and HT antibody staining

Single-cell splenocyte suspensions were isolated as described above and stained with anti-CD45-APC and anti-CD3ε (IGT44) or anti-CD45-APC and gp33/H2-Db, np396/H2-Db, and gp276/H2-Db tetramer in PE (IGT45) and different TotalSeq-C Anti-Mouse Hashtags (TotalSeq-C Anti-Mouse Hashtags 1-10 BioLegend #155861-155879, in staining buffer (DMEM, 2% FCS and 10mM HEPES)) for 20min at 4°C in the dark. Standard spleen cell suspensions were stained under identical conditions but included anti-CD45-FITC (BioLegend 109806, clone 104) to differentiate them from other samples when pooled. Each sample was labeled with a unique hashtag, enabling the downstream assignment of individual cell (10x cell barcode) to their respective source samples. *Cell sorting:* Cells were stained with DAPI just before the sort. For each sample, 500-5000K DAPI^-^, CD45^+^, CD3ε^+^ (IGT44) DAPI^-^, or CD45^+^, tetramer^+^ (IGT45) T cells were sorted on a FACS Aria II and pooled in a single collection tube at 4C. *immgenT TotalSeq-C custom mouse panel staining:* The pooled single cell suspensions were stained with the immgenT TotalSeq-C custom mouse panel, containing 128 antibodies (BioLegend Part no. 900004815) (see NCBI GEO SuperSeries accession number GSE297097 Supplementary Data for complete list) and FcBlock (Bio X Cell #BE0307). Because 500,000 cells are required for staining, unstained splenocytes were spiked in to reach a total of 500,000 cells. *Secondary cell sorting:* Cells were subsequently sorted a second time with the addition of DAPI to select for live T cells and include 5,000 total splenocyte standard cells. Samples were sorted into a single collection tube. See also https://www.immgen.org/ImmGenT/immgenT.SOP.pdf.

### Single-cell RNA and TotalseqC-Sequencing - Library Preparation (immgenT) Cell encapsulation and cDNA library

Single-cell RNA sequencing was performed using the 10x Genomics 5’ v2 platform with Feature Barcoding for Cell Surface Protein and Immune Receptor Mapping, adhering to the manufacturer’s guidelines (CG000330). After cell encapsulation with the Chromium Controller, reverse transcription and PCR amplification were performed in the emulsion. From the amplified cDNA library, smaller fragments containing TotalSeq-C-derived cDNA were separated for Feature Barcode library construction, while larger fragments containing transcript-derived cDNA were preserved for TCR and Gene Expression library generation. Library sizes for both cDNA fractions were evaluated using the Agilent Bioanalyzer 2100 High Sensitivity DNA assay and quantified with a Qubit dsDNA HS Assay kit on a Qubit 4.0 Fluorometer. *RNA library construction:* After enzymatic fragmentation and size selection of the cDNA, the library was ligated to an Illumina R2 sequence and indexed using unique Dual Index TT set A index sequences, SI-TT-B6. *TotalSeq-C library construction:* Totalseq-C derived cDNA was processed into the Feature Barcode libraries following the manufacturer’s protocol. The library was indexed with a unique dual index TN set TN set A (10x part no. 3000510) index sequence SI-TN-F9. *Sequencing:* The three libraries were pooled based on molarity in the following proportions: 47.5% RNA, 47.5% Feature Barcode, and 5% TCR. The pooled libraries were sequenced on an Illumina NovaSeq S2 platform (100 cycles) using the 10x Genomics specifications: 26 cycles for Read 1, 10 cycles for Index 1, 10 cycles for Index 2, and 90 cycles for Read 2.

### Single-cell RNA, TCR and TotalseqC-sequencing - Data processing Code

Code is available on https://github.com/immgen/immgen_t_git/ *Count matrices:* Gene and TotalSeq-C antibody (surface protein panel and hashtags) counts were obtained by aligning reads to the mm10 (GRCm38) mouse genome using the M25 (GRCm38.p6) Gencode annotation and the DNA barcodes for the TotalSeq-C panel. Alignment was performed with CellRanger software (v7.1.0, 10x Genomics) using default parameters. Cells were identified and separated from droplets with high RNA and TotalSeq-C counts by determining inflection points on the total count curve, using the barcodeRanks function from the DropletUtils package. *Sample demultiplexing:* Sample demultiplexing was performed using hashtag counts and the HTODemux function from the Seurat package (Seurat v4.1). Doublets (droplets containing two hashtags) were excluded, and cells were assigned to the hashtag with the highest signal, provided it had at least 10 counts and was more than double the signal of the second most abundant hashtag. Hashtag count data were also visualized using t-SNE to ensure clear separation of clusters corresponding to each hashtag. All single cells from the gene count matrix were uniquely matched to a single hashtag, thereby linking them unambiguously to their original sample. *Quality control (QC) and batch correction:* Cells meeting any of the following criteria were excluded from the analysis: fewer than 500 RNA counts, dead cells with over 10% of counts mapping to mitochondrial genes, fewer than 500 TotalSeq-C counts, or positivity for two isotype controls (indicating non-specific TotalSeq-C antibody binding). Non-T cells were excluded based on the expression of the MNP gene signature, B cell signature, ILC gene signature, and the absence of T cell gene signature (score calculated using UCell::AddModuleScore_UCell). CITEseq data did not meet quality control and was not used in the analysis.

### ImmgenT integration

The data was integrated with the rest of the immgenT dataset using the SCVI.TOTALVI model (v1.2.0) and the 10x lane as a batch parameter. Dimensionality reduction was performed using the pymde.preserve_neighbors() function with default parameters (https://pymde.org/citing/index.html). Cell clustering was carried out using Seurat::FindClusters(). Manual annotation by the immgenT consortium was done using expression of *Cd3, Trbc1,Trbc2, Cd4, Cd8a, Cd8b1, Foxp3, Mki67, Sell, Cd44, Trgc1, Itgax, Itgam*, and *Ms4a1* (RNA) and CD3, TCRB, THY1.2, CD4, CD8A, CD8B, CD62L, CD44, TCRGD, TCRVG1.1, TCRVG2, TCRVG3, CD19, CD20, ITAM.CD11B, ITAX.CD11C, KLRBC-NK1.1 (protein).

### Data accessibility and visualization

Data can be visualized using the Rosetta software https://rosetta.immgen.org/ (IGT44 and IGT45). immgenT data are deposited to NCBI GEO (SuperSeries accession number GSE297097). Specifically, the analyses presented were performed on data fromIGT44 (CD3ε^+^ splenocytes from SPF and NME mice) and IGT45 (LCMV-Arm gp33/H2-D b^+^, np396/H2-Db^+^ or gp276/H2-Db^+^ cells from SPF and NME mice) which can be found in GSEXXXX. More information about ImmgenT can be found on https://www.immgen.org/ImmGenT/.

### Whole blood single-cell RNAseq Data

Blood was collected via the submandibular vein from mice from SPF and NME facilities. After ACK lysis, white blood cells were stained with anti-CD45 (v450) antibody and viability dye (Ghost Dye Red 780) prior to FACS (BD FACSAria II) purification of live, CD45+ cells at the University of Minnesota Flow Cytometry Core. Encapsulation, library preparation, and library sequencing were performed by the University of Minnesota Genomics Center, using the 10x Chromium Controller, 10x Genomics Chromium Single Cell 3′ Reagent Kits (v2, CG00052), and Illumina NovaSeq 6000. Resulting reads were aligned to the mm10 (GRCm38) mouse genome with the 10x Genomics CellRanger software (v3.0.2). Cells with fewer than 500 or greater than 2,500 unique genes and those with mitochondrial genes accounting for >5% of total counts were excluded from downstream analyses. Count normalization and variance stabilization was performed on output from each sample with sctransform::SCTransform() (v2) in R using 3000 variable features, after which PCA, neighbor selection, clustering (res = 0.6), and UMAP dimensionality reductions were performed using PCA dimensions 1:30. Normalized data from each sample were then integrated using Seurat::IntegrateLayers() using the RPCA integration method. Neighbor selection, clustering (res = 0.6), and UMAP dimensionality reductions were performed on the integrated data using RPCA dimensions 1:30.

### Differential Gene Expression (DEG) Analysis

All differential expression analyses were performed using Seurat::FindMarkers() (v5.3.0). DEGs were defined as genes expressed in >25% of cells from either sample group, >1 Log2 average Fold Change between sample groups, and an adjusted p value < .05.

### Cluster Representation Analysis

Differential cluster representation was determined using speckle::propeller() (v3.2.1). To assess differences in cluster representation across sample groups, subclusters were first grouped by cell identity prior to the appropriate statistical test (ANOVA, t-test). Significant compositional differences were defined at an FDR cutoff of .05.

### Module scores

Module scoring was performed using UCell::AddModuleScore_UCell() (v2.10.1) in R. The following gene lists were used to generate signatures: LLEC module score list– *‘Prf1’, ‘Gzmb’, ‘Gzma’, ‘Klrc3’, ‘Klra9’, ‘Klrb1c’, ‘Klrc1’, ‘Klre1’, ‘Klrg1’, ‘Ccl4’, ‘Ccl9’, ‘Cx3cr1’, ‘S1pr5’, ‘Runx1’, ‘Bhlhe40’, ‘Zeb2’, ‘Prdm1’*; TCM module score list– *‘Sell’, ‘Il7r’, ‘Cxcr5’, ‘Cxcr3’, ‘Slamf6’, ‘Ccr7’, ‘Itgax’, ‘Eomes’, ‘Myc’, ‘Id3’, ‘Tcf7’*.

### Percent Expression

The percent of cells within conditions expressing indicated genes was determined using scCustomize::Percent_Expressing() (v3.2.0).

### Visualizations

Volcano plots were generated using the EnhancedVolcano (v1.24.0) package in R. Bar plots were generated using dittoSeq::dittoBarPlot() (v1.18.0) in R. Gene density plots were generated using Nebulosa::plot_density() (v1.18.0) in R. immgenT minimal distortion embeddings were added using ZemmourLib::AddLatentData() (https://github.com/dzemmour/ZemmourLib). All other scRNA-sequencing visualizations were made with native Seurat plotting functions.

## Data Availability

immgenT scRNA-sequencing data have been deposited to NCBI GEO (SuperSeries accession number GSE297097). Specifically, the analyses presented were performed on IGT44 (CD3ε^+^ splenocytes from SPF and NME mice) and IGT45 (LCMV-Arm gp33/H2-D^b+^, np396/H2-D^b+^ or gp276/H2-D^b+^ cells from SPF and NME mice). For Figure 1, additional SPF splenocyte samples from IGT10, 36, and 40 were included as biological replicates. See all immgenT clusters and maps here: https://rstats.immgen.org/immgenT_interactive_maps/CD8.html Figure 2 scRNA-seq data have been deposited to NCBI GEO (Series accession number GSE310286). For code used in the sequencing analyses see: https://GitHub.com/thefa002/nme_igt

## Acknowledgements

We thank the NIH Tetramer Core Facility (NIH Contract 75N93020D00005 and RRID:SCR 026557) for providing monomers. We thank the University of Minnesota Masonic Cancer Center Flow Cytometry Resource, Research Animal Resources, and BSL3 facility management for their expertise and support of this work.

## Collaborators

Participants in the immgenT Project include:

Aaron Liu^1^, Alexander Chervonsky^2^, Alexandra Cassano^2^, Alia Welsh^3^, Amir Ferry^11^, Ananda Goldrath^11^, Andrea Lebron-Figueroa^5^, Ankit Malik^2^, Anna-Maria Globig^4^, Antoine Freuchet^2^, Bana Jabri^2^, Charlotte Imianowski^6^, Christophe Benoist^5^, Claire Thefaine^7^, Dan Kaplan^6^, Dania Mallah^5^, Dario Vignali^6^, David Sinclair^5^, David Zemmour^2^, Derek Bangs^8^, Domenic Abbondanza^2^, Enxhi Ferraj^9^, Eric Weiss^6^, Erin Lucas^7^, Evelyn Chang^9^, Gavyn Chern Wei Bee^10^, Giovanni Galletti^11^, Ian Magill^5^, Iliyan D Iliev^12^, Joonsoo Kang^9^, Jordan Voisine^2^, Josh Choi^5^, Julia Merkenschlager^13^, Jun R. Huh^5^, Katharine Block^7^, Ken Cadwell^10^, Kennidy K. Takehara^11^, Kevin Osum^7^, Laurent Brossay^14^, Laurent Gapin^15^, Liang Yang^5^, Lizzie Garcia-Rivera^1^, Marc K. Jenkins^7^, Maria Brbic^16^, Maria-Luisa Alegre^2^, Marion Pepper^8^, Mariya London^17^, Matthew Stephens^2^, Maurizio Fiusco^16^, Melanie Vacchio^3^, Michael Starnbach^5^, Michel Nussenzweig^13^, Mitch Kronenberg^18^, Myriam Croze^19^, Nalat Siwapornchai^5^, Nathan Morris^12^, Nicole E. Scharping^11^, Nika Abdollahi^19^, Nitya Mehrotra^2^, Odhran Casey^5^, Olga Barreiro del Rio^5^, Paul Thomas^20^, Peter Carbonetto^2^, Remy Bosselut^3^, Rocky Lai^9^, Sam Behar^9^, Sam Borys^14^, Sara E. Hamilton^7^, Sara Mostafavi^8^, Sara Quon^11^, Serge Candéias^21^, Shanelle Reilly^14^, Shanshan Zhang^5^, Siba Smarak Panigrahi^16^, Sofia Kossida^19^, Stefan Muljo^3^, Stefan Schattgen^20^, Stefani Spranger^22^, Steve Jameson^7^, Susan M. Kaech^1^, Takato Kusakabe^12^, Taylor Heim^22^, Tianze Wang^8^, Tomoyo Shinkawa^9^, Ulrich von Andrian^5^, Val Piekarsa^5^, Véronique Giudicelli^19^, Vijay Kuchroo^5^, Woan-Yu Lin^12^, Ziang Zhang^2^

1. NOMIS Center, Salk Institute for Biological Sciences, 2. The University of Chicago, 3. National Institutes of Health, 4. Allen Institute for Immunology, 5. Harvard Medical School, 6. Dept of Dermatology and Immunology, University of Pittsburgh, 7. University of Minnesota, 8. University of Washington, 9. UMass Chan Medical School, 10. University of Pennsylvania, 11. University of California San Diego, 12. Weill Cornell Medicine, 13. The Rockefeller University, 14. Brown University, 15. University of Colorado Anschutz Medical Campus, 16. Swiss Federal Institute of Technology, Lausanne, 17. New York University, 18. La Jolla Institute, 19. IMGT, Univ Montpellier, 20. St. Jude Children’s Research Hospital, 21. Alternative Energies and Atomic Energy Commission, Grenoble, 22. Massachusetts Institute of Technology

