## Supplementary Figures for "Diverse Microbial Exposure Enhances CD8^+^ T Cell Effector Memory Output and Function"

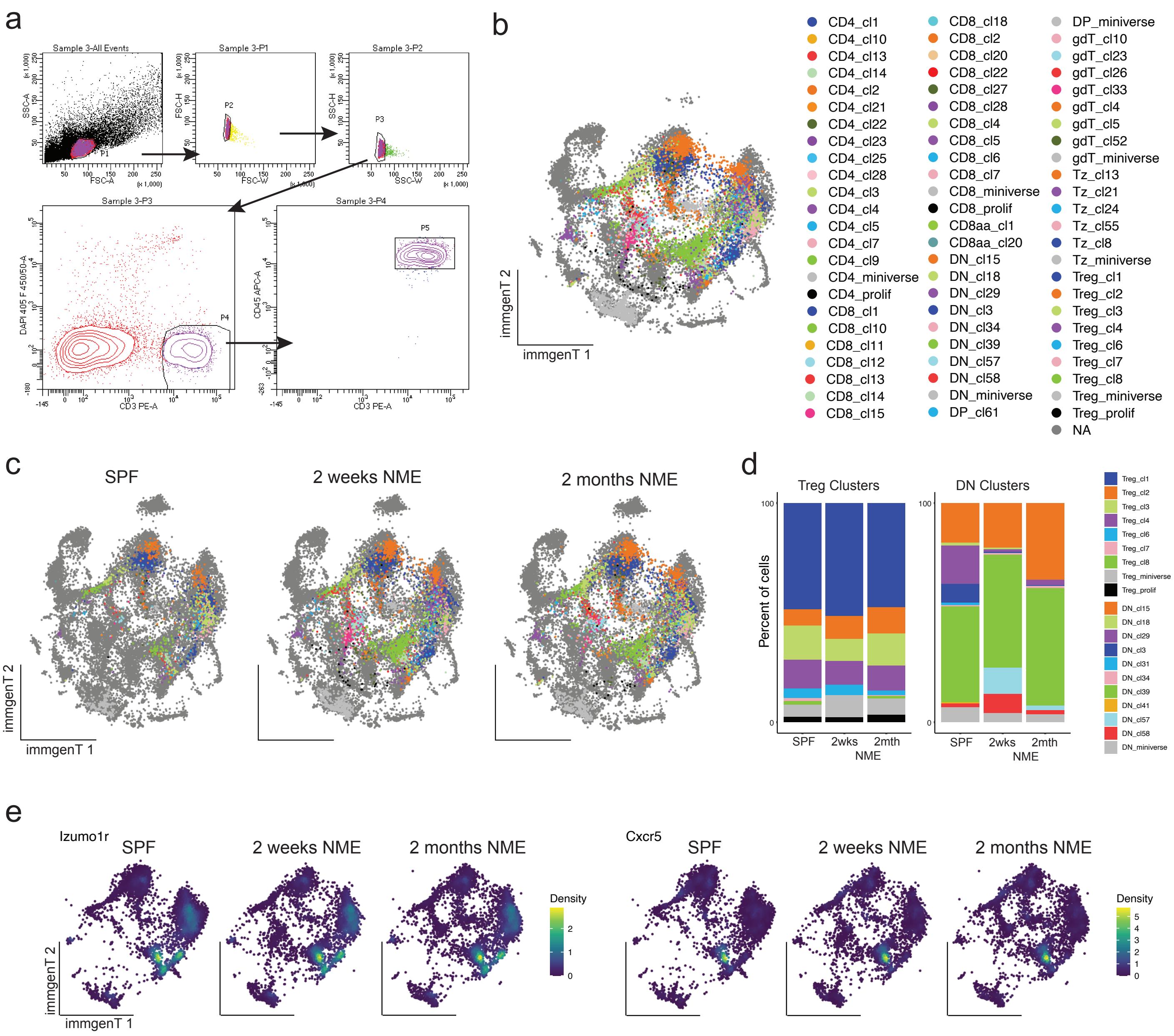
a
Sample 3-All Events
Sample 3-P1
Sample 3-P2
Sample 3-P3
Sample 3-P4
SSC-A
FSC-A
FSC-H
FSC-W
SSC-H
SSC-W
P1
P2
P3
P4
P5
DAPI 405 F 450/50-A
CD3 PE-A
CD45 APC-A
b
immgenT 1
immgenT 2
CD4_cl1
CD4_cl10
CD4_cl13
CD4_cl14
CD4_cl2
CD4_cl21
CD4_cl22
CD4_cl23
CD4_cl25
CD4_cl28
CD4_cl3
CD4_cl4
CD4_cl5
CD4_cl7
CD4_cl9
CD4_miniverse
CD4_prolif
CD8_cl1
CD8_cl10
CD8_cl11
CD8_cl12
CD8_cl13
CD8_cl14
CD8_cl15
CD8_cl18
CD8_cl2
CD8_cl20
CD8_cl22
CD8_cl27
CD8_cl28
CD8_cl4
CD8_cl5
CD8_cl6
CD8_cl7
CD8_miniverse
CD8_prolif
CD8aa_cl1
CD8aa_cl20
DN_cl15
DN_cl18
DN_cl29
DN_cl3
DN_cl34
DN_cl39
DN_cl57
DN_cl58
DN_miniverse
DP_cl61
DP_miniverse
gdT_cl10
gdT_cl23
gdT_cl26
gdT_cl33
gdT_cl4
gdT_cl5
gdT_cl52
gdT_miniverse
Tz_cl13
Tz_cl21
Tz_cl24
Tz_cl55
Tz_cl8
Tz_miniverse
Treg_cl1
Treg_cl2
Treg_cl3
Treg_cl4
Treg_cl6
Treg_cl7
Treg_cl8
Treg_miniverse
Treg_prolif
NA
c
SPF
2 weeks NME
2 months NME
immgenT 1
immgenT 2
d
Treg Clusters
DN Clusters
Percent of cells
100
0
SPF
2wks
2mth
NME
Treg_cl1
Treg_cl2
Treg_cl3
Treg_cl4
Treg_cl6
Treg_cl7
Treg_cl8
Treg_miniverse
Treg_prolif
DN_cl15
DN_cl18
DN_cl29
DN_cl3
DN_cl31
DN_cl34
DN_cl39
DN_cl41
DN_cl57
DN_cl58
DN_miniverse
e
Izumo1r
SPF
2 weeks NME
2 months NME
Density
Cxcr5
immgenT 1
immgenT 2

**Supplementary Figure 1:** a) Gating strategy for initial sort of CD3ϵ$^{+}$ cells for scRNA sequencing. b) immgenT dimensions of CD3ϵ$^{+}$ splenocytes from SPF and NME mice followed 2 weeks or 2 months of cohousing (n = 4 each) overlaid on the complete immgenT atlas. c) immgenT dimensions, split by condition (SPF n= 1, NME n = 4 each). d) Frequency of immgenT Treg and DN T cell subclusters by condition. e) Expression levels of indicated genes in SPF and NME T cells.

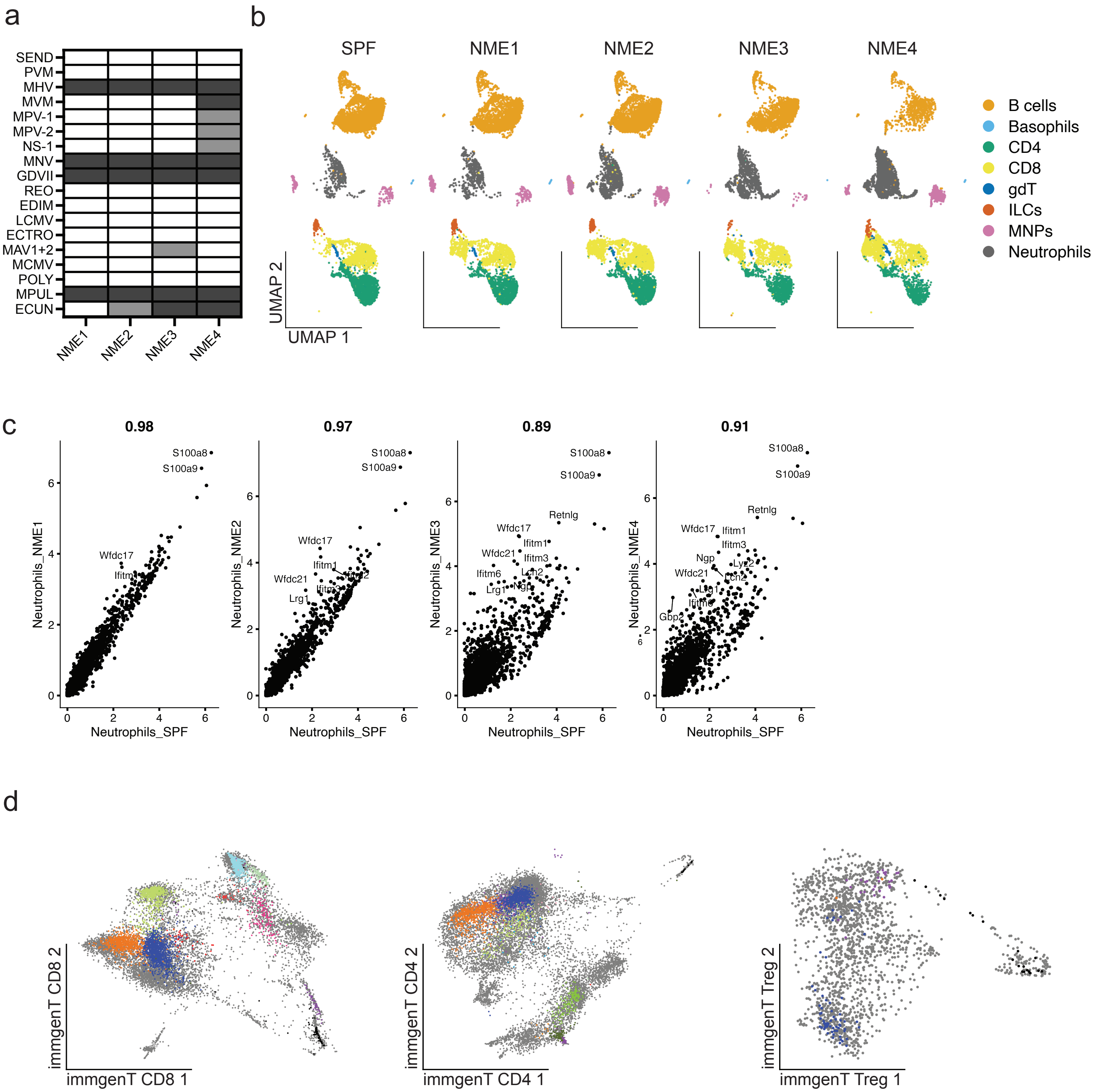
a
SEND
PVM
MHV
MVM
MPV-1
MPV-2
NS-1
MNV
GDVII
REO
EDIM
LCMV
ECTRO
MAV1+2
MCMV
POLY
MPUL
ECUN
NME1
NME2
NME3
NME4
b
SPF
NME1
NME2
NME3
NME4
B cells
Basophils
CD4
CD8
gdT
ILCs
MNPs
Neutrophils
UMAP 2
UMAP 1
c
0.98
0.97
0.89
0.91
S100a8
S100a9
Wfdc17
Wfdc21
Ifitm1
Ifitm3
Ifitm6
Lrg1
Retnlg
Ngp
Lcn2
Ly6g
Gbp2
Neutrophils_NME1
Neutrophils_NME2
Neutrophils_NME3
Neutrophils_NME4
Neutrophils_SPF
d
immgenT CD8 2
immgenT CD8 1
immgenT CD4 2
immgenT CD4 1
immgenT Treg 2
immgenT Treg 1

**Supplementary Figure 2:** a) Serology results from NME mice following 60 days of cohousing. Dark gray = moderate to strong positive, light gray = equivocal (weak positive), white = negative. SEND = murine respirovirus (Sendai virus), PVM = pneumonia virus of mice, MHV = mouse hepatitis virus (mouse coronavirus), MVM = minute virus of mice, MPV-1 = mouse parvovirus type 1, MPV-2 = mouse parvovirus type 2, NS-1 = nonstructural protein 1 of parvoviruses, MNV = murine norovirus, GDVII = Theiler's murine encephalomyelitis virus (mouse theilovirus), REO = reovirus, EDIM = rotavirus, LCMV = lymphocytic choriomeningitis virus, ECTRO = Ectromelia (mousepox), MAV1+2 = mouse adenovirus 1 and 2, MCMV = murine cytomegalovirus, POLY = mouse polyomavirus, MPUL = *Mycoplasma pulmonis*, ECUN =*Encephalitozoon cuniculi*. b) Integrated UMAP clustering as in Fig 2B, split by condition. c) Comparison of aggregated neutrophil gene expression between SPF and NME samples. d) Integrated UMAP clustering of either CD8, CD4 or Treg T cells.

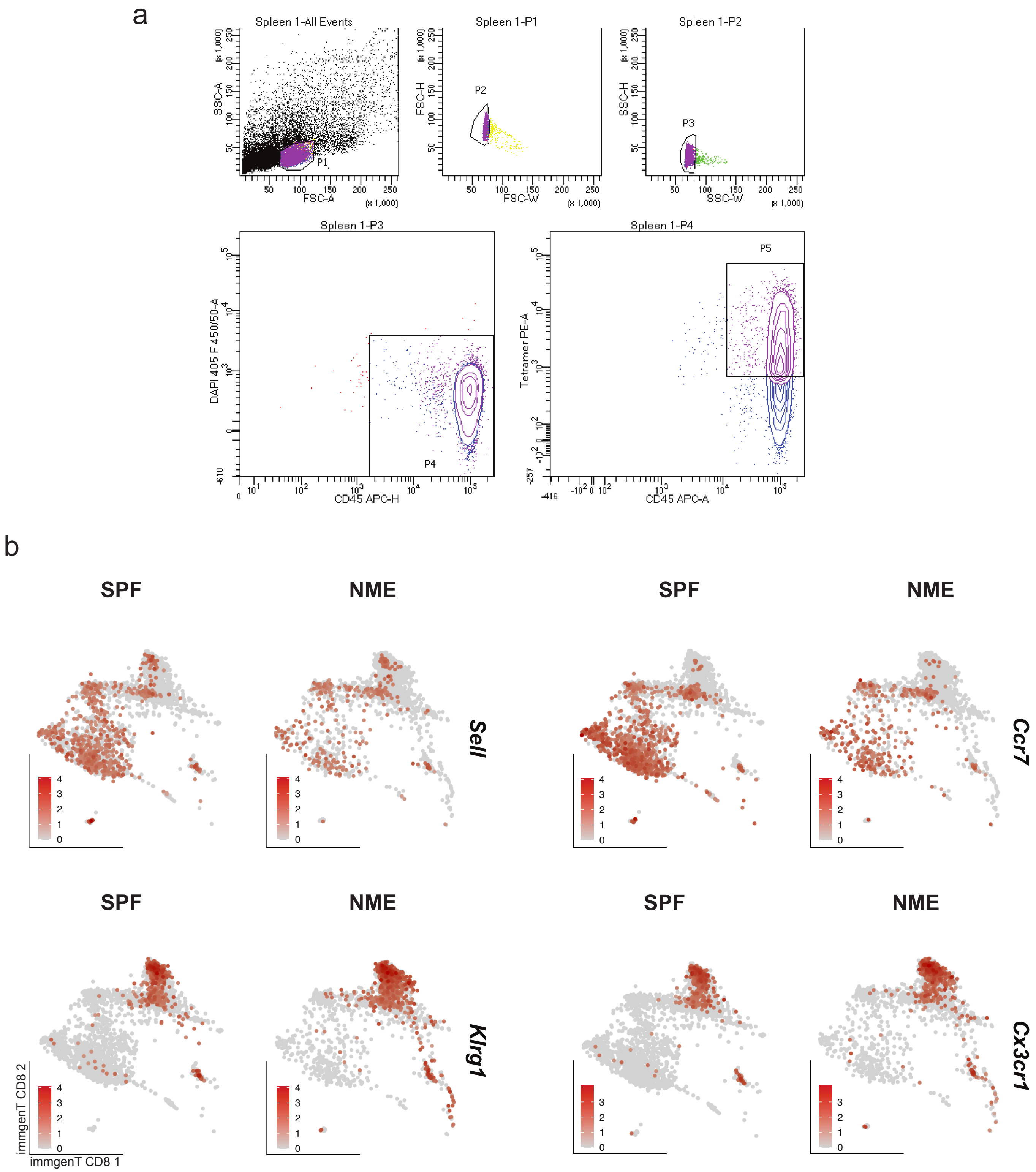


**Supplementary Figure 3:** a) Gating strategy for initial sort of gp33/H2-D$^{b}$, np396H2-D$^{b}$ or gp276/H2-D$^{b}$ tetramer$^{+}$ cells for scRNA-sequencing. b) Expression of indicated genes by SPF or NME CD8 T cells from Figure 3.

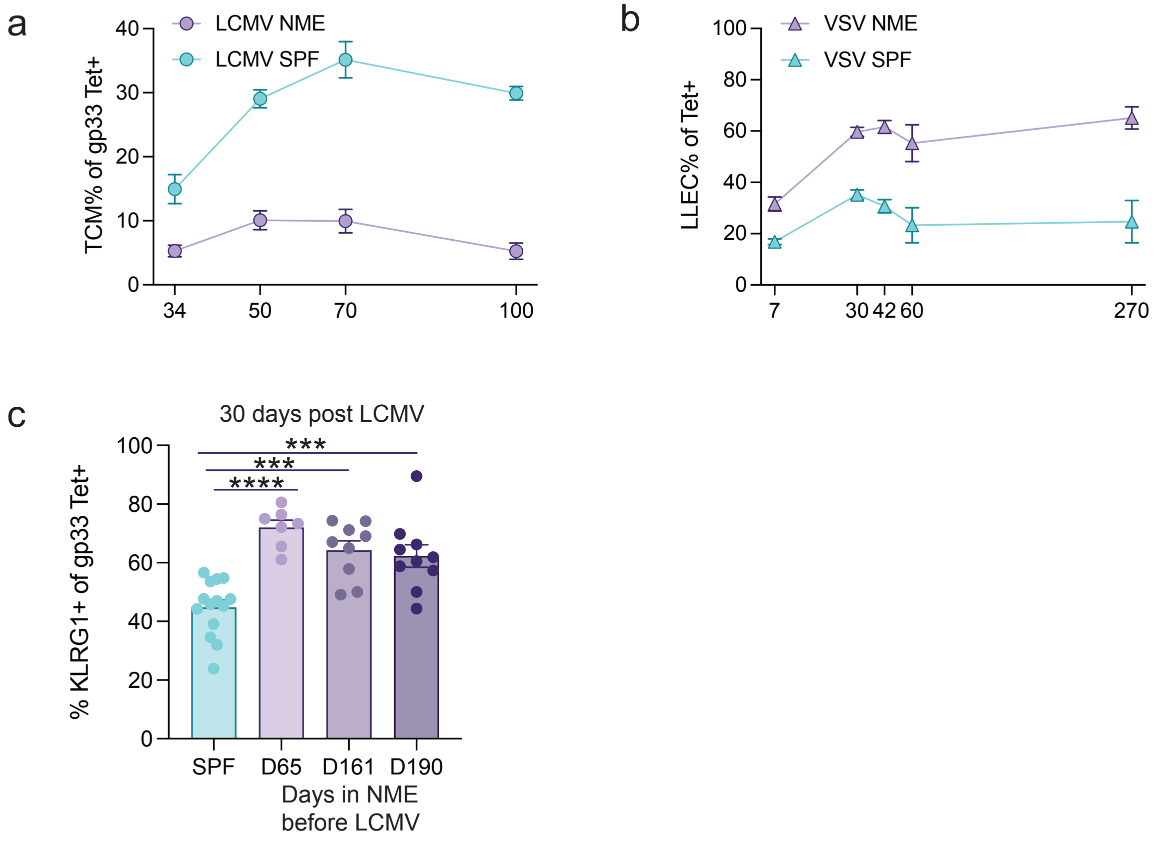


**Supplementary Figure 4:** a) Frequency of gp33 tetramer$^{+}$ MPEC/TCM (CD62L+) in the blood of mice from Fig 3 over time. b) Frequency of KLRG1$^{+}$, SIINFEKL tetramer$^{+}$, CD8$^{+}$ T cells in the blood over time following infection with VSV-OVA. c) Frequency of KLRG1$^{+}$ gp33 tetramer$^{+}$ CD8$^{+}$ T cells in the blood of SPF and NME mice 30 d.p.i. with LCMV-Arm. For c a 1-way Anova followed by Tukey's multiple comparisons test was used.

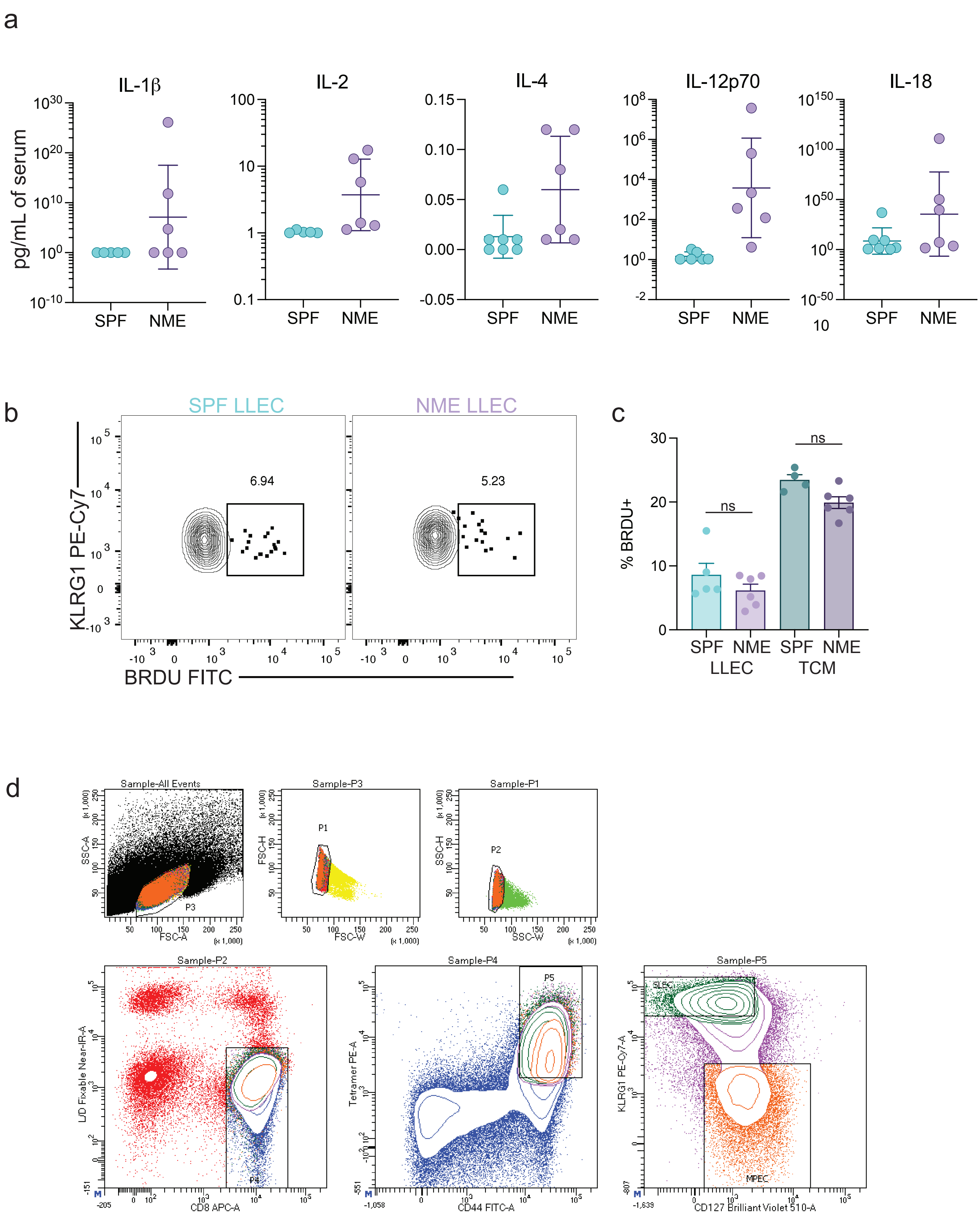
a
IL-1β
IL-2
IL-4
IL-12p70
IL-18
pg/mL of serum
SPF
NME
b
SPF LLEC
NME LLEC
6.94
5.23
KLRG1 PE-Cy7
BRDU FITC
c
% BRDU+
ns
SPF NME SPF NME
LLEC TCM
d
Sample-All Events
Sample-P3
Sample-P1
P3
P1
P2
SSC-A
FSC-H
SSC-H
FSC-A
FSC-W
SSC-W
Sample-P2
Sample-P4
Sample-P5
P4
P5
SLEC
MPEC
L/D Fixable Near-IR-A
CD8 APC-A
Tetramer PE-A
CD44 FITC-A
KLRG1 PE-Cy7-A
CD127 Brilliant Violet 510-A

**Supplementary Figure 5:** a) Quantification of serum cytokine levels from SPF and NME mice. NME mice were cohoused for 150-200 days prior to serum collection. b) Representative flow cytometry plots showing BrdU incorporation by gp33 tetramer$^{+}$ LLECs in SPF and NME mice, quantified in c. c) Frequency of BrdU incorporation by gp33 tetramer$^{+}$ TCM and LLEC. d) Sorting strategy for SLEC and MPECs from SPF mice. Related to Figure 5a-c.
